# Agile workflow for interactive analysis of mass cytometry data

**DOI:** 10.1101/2020.05.28.120527

**Authors:** Julia Casado, Oskari Lehtonen, Ville Rantanen, Katja Kaipio, Luca Pasquini, Antti Häkkinen, Elenora Petrucci, Olli Carpén, Mauro Biffoni, Anniina Färkkilä, Sampsa Hautaniemi

## Abstract

**Motivation:** Single-cell proteomics technologies, such as mass cytometry, have enabled characterization of cell-to-cell variation and cell populations at a single cell resolution. These large amounts of data, however, require dedicated, interactive tools for translating the data into knowledge.

**Results:** We present a comprehensive, interactive method called *Cyto* to streamline analysis of large-scale cytometry data. *Cyto* is a workflow-based open-source solution that automatizes the use of of state-of-the-art single-cell analysis methods with interactive visualization. We show the utility of *Cyto* by applying it to mass cytometry data from peripheral blood and high-grade serous ovarian cancer (HGSOC) samples. Our results show that Cyto is able to reliably capture the immune cell sub-populations from peripheral blood as well as cellular compositions of unique immune- and cancer cell subpopulations in HGSOC tumor and ascites samples.

**Availability:** The method is available as a Docker container at https://hub.docker.com/r/anduril/cyto and the user guide and source code are available at https://bitbucket.org/anduril-dev/cyto

**Supplementary information:** Supplementary material is available and FCS files are hosted at flowrepository.org/id/FR-FCM-Z2LW

## 1 Introduction

Single-cell technologies such as Cytometry by Time-Of-Flight (CyTOF), multiplexed imaging, or single cell RNA sequencing have enabled characterizating tumor-microenvironment compositions and cell populations at a single-cell resolution (Galli et al., 2019). However, currently the pace at which insight is extracted from massive single-cell data sets remains the same as with the previous low-throughput technologies (Brodin, 2018). Common CyTOF analysis steps have steadily reached a quasi-standard workflow that involves manual gating with FlowJo™ or other 2D scatter plot tools followed by dimensionality reduction with t-SNE (Van Der Maaten and Hinton, 2008) and unsupervised clustering. Typically these analyses are executed with different software or platforms, which maes the resutls prone to errors and biases. Meanwhile, each new experiment requires a new set of custom scripts to fit the analysis needs, and new computational methods and algorithms are being developed at a fast rate (Qiu, 2017; Angerer et al., 2016; Höllt et al., 2016). The most comprehensive semiautomatic workflow available is CytoBank (Kotecha et al., 2010), a commercially available service that allows the users to load the data to a cloud and perform analyses without the need for advanced technical skills. Open-source alternatives have been developed to make analysis accessible. For example, Cytofkit (Chen et al., 2016), integrate methods available only within the R ecosystem and no parallelization support due to R limitations, which is suboptimal when analyzing very large data sets. Other, more complex solutions, such as Cytosplore (van Unen et al., 2017) and CYT (Amir et al., 2013), allow for only one method for each step of the analysis, one transformation type, one sampling approach, one clustering algorithm, and one dimensionality reduction method. Furtheremore, none of these software does not support iterative analysis, which is required for rapidly testing hypotheses and ideas during analysis. Iterative analysis is recognized as a key requirement for workflow languages (Almeida, 2010), and it is particularly important in the analysis of mass cytometry data as the data sets are complex and require testing different parameter settings, algorithms, etc. in an iterative and interactive fashion. We have designed and implemented an analysis software *Cyto* that enables interactive analysis and meets the need for accessibility to and reporting of reproducible methods.

We demonstrate the utility of *Cyto* with two CyTOF datasets. Firstly, we use control data from peripheral blood mononuclear cells (PBMC) (Van Unen et al., 2016) to demonstrate fast quality assessment of the data and recapitulation of the previous findings in only two iterations of analysis. Secondly, we applied *Cyto* on a dataset from high-grade serous ovarian cancer (HGSOC). By applying *Cyto* on this dataset, we were able to rapidly measure abundance of cell types, and single-out specific tumor cell populations facilitating biological discovery and clinical interpretation of high dimensional single-cell cytometry data.

## 2 System and Methods

Cyto is built on top of the workflow framework Anduril 2 (Cervera et al., 2019), a language-agnostic framework that enables rapid integration of new and old methods as building blocks.

### 2.1 Cyto modules

#### 2.1.1 Graphical user interface

The user interface was developed as a light Flask application server within a Docker container. Docker avoids dependency installation and versioning issues, and therefore eases compatibility between researchers. The application handles data upload and download and saves user configuration changes. All projects are saved locally in the user’s computer in case Docker is restarted.

#### 2.1.2 Interactive results browser

To make *Cyto* modular, the user-data interaction was implemented as a separate web application built with Python dashboards, a powerful framework that supports interactive Plotly components. The choice of visualization strategies are based on those reported in relevant publications, particularly in Nowicka *et al.*, 2017.

#### 2.1.3 Cytometry analysis pipeline

The analysis pipeline is shown in Supplementary Figure S1. Briefly, integration of cytometry specific methods was achieved through development of new Anduril components built with MATLAB^®^, R, Python, Java, or Bash scripts, depending on the programming language of the original implementation of each method. A list of the currently integrated methods for data processing, clustering, 2D embedding, and building dashboard are listed in Supplementary Table S1.

### 2.2 Materials and methods for peripheral blood case study

#### 2.2.1 Data acquisition of peripheral blood myeloid cells

We downloaded the mass cytometry FCS files from (Van Unen *et al.*, 2016) and selected the control (Ctrl) samples (*n*=14) to recapitulate the PBMC cell subtypes. No preprocessing of the data was required before the *Cyto* analysis.

#### 2.2.2 Cyto analysis of data quality

We selected the channels used in (Van Unen *et al.*, 2016). The complete dataset contained 48,611,486 cells, of which we randomly subsampled to 300,000 cells and transformed all selected channels with an arcsinh transformation (cofactor 5). The parameters and their values are listed in Supplementary File S1. Multidimensional scaling (MDS) and non-redundancy scores (NRS) visualization within *Cyto* Dash report were used to identify outlier samples.

#### 2.2.3 *Cyto* recapitulation of cell types

After excluding the outlier samples 52_CtrlAdult5_PBMC and 53_CtrlAdult6_PBMC we ran *Cyto* analysis (Supplementary File S2) on the remaining 12 Ctrl samples. This dataset contained 41,779,615 cells which were randomly downsampled to 300,000 cells. The same parameters as in the previous iteration were used but clustering was done with FlowSOM algorithm (*k*=18) and dimensionality reduction by tSNE (*n*=10,000; perplexity=20; theta=0.3). The cell type labels used and prior knowledge of marker expression profiles are described in (Van Unen *et al.*, 2016).

### 2.3 Materials and methods for case study HGSOC case study

#### 2.3.1 Data acquisition of High-Grade Serous Ovarian Cancer

Tissue and ascites specimens were collected from 15 consented patients (Supplementary Table S2) at the Department of Obstetrics and Gynecology, Turku University Central Hospital. Samples were analysed with CyTOF 1 mass cytometer (DVS Sciences Fluidigm). The antibody panel was manually curated with focus on markers of cell populations that compose the tumor compartment and less attention to the microenvironment (Supplementary Table S2). For further details about sample preparation and CyTOF assay see Supplementary Methods.

#### 2.3.2 Tumor compartment identification with *Cyto*

The FCS files and the CSV file with clinical annotations were uploaded to *Cyto* and processed as shown in Supplementary File S3. 300,000 cells were randomly sampled from a total of 65,331,333 cells in the complete dataset. After the cyto run with signal transformation *log1p*, sample-wise mean centering, clustering with Phenograph (*k*=200), and dimensionality reduction by UMAP (*n*=10,000, min-dist=0.1, knn=90). we associate cell types to each cluster based on the expression of canonical cell type markers (Supplementary Figure S5). The clustering results were downloaded from *Cyto* to label the clusters and compare global cell type abundances. To maximize the number of tumor cells we ran a second iteration of analysis using a density-biased downsampling while keeping all other parameters unchanged (Supplementary Figure S6). The resulting CSV file was filtered in AWK to keep only the tumor cells for the next iteration.

#### 2.3.3 Tumor cell population analysis

All tumor cells were used with no preprocessing (setting *none*). We applied all clustering methods to show the different effect of complex cell populations that do not follow a clear lineage on clustering results, each analysis is detailed with the method name within the configuration file Supplementary File S4.

## 3 Results

Cyto is an open-source application that enables running cytometry analysis pipelines that integrate state-of-the-art tools with reliable reporting and reproducibility as shown in Figure 1. Importantly, the interactive visualization of the results removes the need for many iterations of editing the common analysis scripts and improving interpretation time over traditional static visualization methods (Dix and Ellis, 1998).

**Fig 1.**
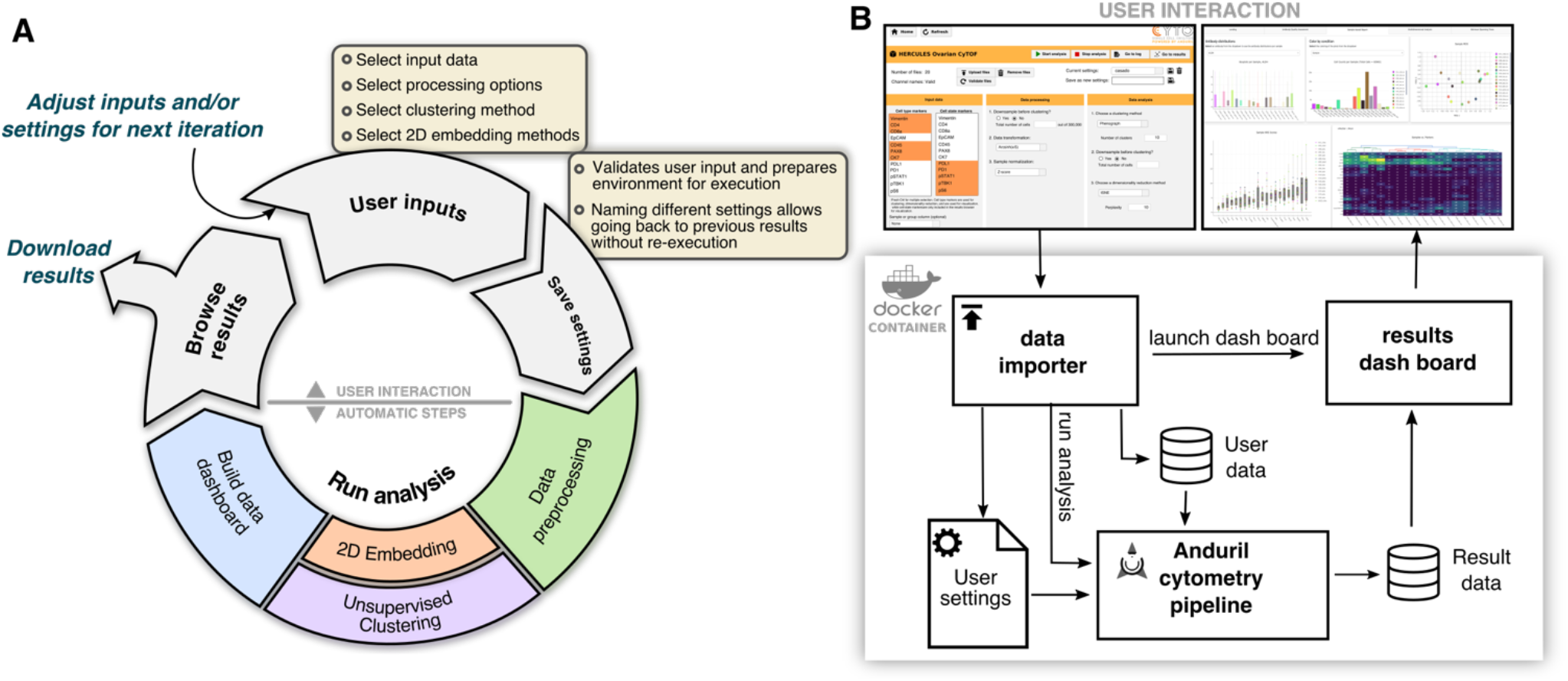
Workflow for cytometry analyses. (A) Diagram of steps showing cytometry analysis as an iterative process and how our framework enables knowledge discovery. (B) Schematic of the analysis environment to enable multi-system compatibility. On top screenshots of the *data importer* and the *results browser* as the two separate python applications.

### 3.1 Software architecture supports reproducibility and accessibility requirements

The design of *Cyto* was driven by both the need of iterative analysis characteristic to the high dimensional cytometry field and the requirement for easily reporting of methods and parameters used in each step of an analysis, which are critical for reproducibility. With this in mind, we developed Anduril components to integrate the most popular cytometry tools into fully customizable analysis pipelines (https://bitbucket.org/anduril-dev/cytometry and https://bitbucket.org/anduril-dev/tools).

Our cytometry analysis pipeline includes tools from different fields and in different languages that are wrapped into modular units, called components, which are interchangeable and reusable throughout the pipeline development process. To enable rapid changes to the choice of components and to support non-bioinformaticians to interact with CyTOF datasets, we built a lightweight user interface that runs a generalizable Anduril pipeline (Supplementary Figure S1). This is achieved with two web-based Python applications: the first one is the *data importer* where the user defines their analysis parameters, while the second one is the *results browser* to enable interactive data visualization through Plotly figures. Finally, to simplify installation requirements and thus enhance accesibility, we packaged this system into an interactive Docker containter which can run on most operating systems. To our knowledge, *Cyto* is the first open-source solution that features access to multiple cytometry tools with a low learning threshold for non-bioinfomaticians.

#### 3.1.1 Mass cytometry data analysis with Cyto

On a general scale, *Cyto* follows a common CyTOF workflow (Figure 1A and Supplementary Figure S1), however, each step enables agile and fast iterations. The *preprocessing* components are a critical step of a CyTOF pipeline. An *arcsinh* transformation is usually applied and it works well in many experiments, however, it may truncate high values to an artificial maximum. For this reason, users may choose also logarithmic or quadratic scaling. Other important parts of the preprocessing implemented in new components are quality assessment, normalization, gating, and filtering components. By generalizing these steps in the *Cyto* pipeline instead of running multiple independent scripts or manual analysis, the user has a comprehensive log of methods tested, and complete control of the preprocessing steps without having to code all the logic that is already included in each component.

Because of the flexibility to adapt new tools as components to this bundle, *Cyto* supports *dimensionality reduction* and *unsupervised clustering* methods, along with new tools that can be included when available. The third popular toolbox contains *lineage inference* methods; we integrated them to produce an output that can be further analysed with any component or visualized with the *interactive visualization* components. The interactive visualization components transform data into plotly objects to be used either locally in the user’s browser or included in a Dash application, as demonstrated in the *Cyto* method. Lastly, Anduril counts with a large *tools* bundle with components for statistical analysis, CSV file manipulations, and machine learning analysis, all of which are fully compatible with our cytometry components.

#### 3.1.2 Cyto design enables customized analysis steps

Figure 1A depicts worfklow for a standard cytometry analysis project. First, the user sets the input data and parameters for the analysis in the *data importer*. Different types of research questions require different settings. Questions about population abundance can analyze all cells or a random sample, while detection and identification of rare cell populations requires a density-biased sample as implemented in SPADE package (Qiu *et al.*, 2011) to preserve smaller populations. Commonly used clustering algorithms in the field are tailored for different research setups (Weber and Robinson, 2016). Algorithms based on a k-nearest neighbors approach are suitable for samples where the expression of markers varies smoothly, e.g. are expected to belong to an evolving. On the other hand, samples with distant subpopulations will benefit from a more fragmented clustering method, like k-means. Thus, it is important to support the use of the right tool for the right question, not just the easiest to use. Second, the user saves the settings. At the moment of saving these options, *Cyto* validates the inputs and creates a new execution folder, which is used to archive the configuration, to support reproducibility, and to store the intermediate results, to support re-running only necessary steps on following iterations. Third, starting the analysis will launch the cytometry analysis pipeline and build the *results browser*. Upon completion of the analysis, the browser will enable the user to build new hypotheses and make informed decisions for the next iterations. The browser helps interacting with high dimensional data and multiple results effectively, from assessing signal quality and sample selection quality to examining individual or groups of cell populations. In the *data importer*, we can also download the results as a table that includes all preprocessed data and clustering results, and the *results browser* can also be downloaded to be hosted on a web server as supporting material for complex publication results.

The presented cytometry components can also be integrated into Anduril pipelines independently of our proposed analysis pipeline within the *Cyto* system. Independent pipelines are specially useful for laboratories with highly specific research questions that cannot be addressed within the *Cyto* system but benefit from some of the steps. The modular design of our method enables other researchers to follow this design for specialized needs (Figure 1B and Supplementary Figure S1).

### 3.2 Case study I: Peripheral Blood Myeloid Cells dataset

#### 3.2.1 Interactive browser enables outlier detection

The *results browser* generates summary figures to assess data quality. Multidimensional scaling visualization of the average expresion on each sample (Figure 2A) highlights sample *53_CtrlAdult6_PBMC* as an outlier at the general level. While visualization of Non-Redundancy Scores (Figure 2B), identifies also sample *52_CtrlAdult5_PBMC* due to artifactually low signal, seen as lowest NRS for more than 50% of the antibodies. Further assessment of outlier samples is possible by exploring the profiles of cell populations predominant in the outlier population (Supplementary Figure S2). In this analysis, sample *53_CtrlAdult6_PBMC* shows over-representation of myeloid cells, possibly caused by preanalytical conditions. Sample *52_CtrlAdult5_PBMC* shows a very low Simpson’s diversity index (0.34) compared with the rest of the samples (μ=0.67; σ=0.003) (Supplementary Figure S3). By creating a new analysis from the *data importer*, we were able to rapidly discard poor quality samples and repeat the analysis with the same settings.

**Fig 2.**
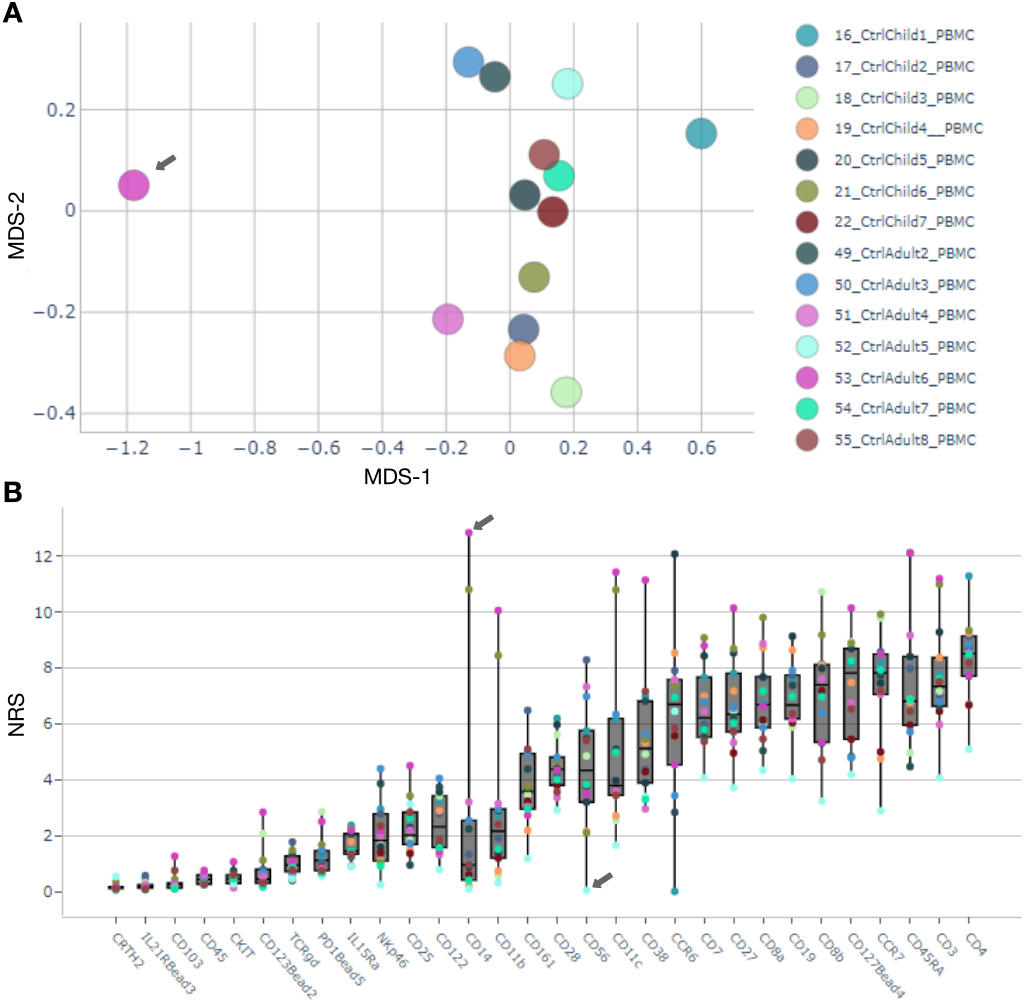
Easy outlier detection and characterization. (A) MDS plot shows sample 53_CtrlAdult6_PBMC separate from the other Ctrl samples. (B) Non-redundancy scores visualization; sample 53_CtrlAdult6_PBMC has highest NRS on marker CD14, and sample 52_CtrlAdult5_PBMC shows lowest for 18 out of 30 markers.

#### 3.2.2 Cyto recapitulates cell-type identification from PBMCs

We set out to test the performance of *Cyto* in detecting immune cell populations from the PBMC dataset. By using density-biased sampling, we quickly recapitulate the cell types present in these samples in line with the authors of the data. Figure 3 shows the results from *Cyto* manually colored by the cell type classification for each cell. Visual separation of some cell types can be further explored by intensity tSNEs and lineage trees (Supplementary Figure S4). Interactive visualization of relevant markers shows slight differences in expression within the same cell type. Additionally, the lineages presented as the minimum spanning trees can be applied to the result of any clustering algorithm. *Cyto* analysis workflow herein reliably identifies biological cell populations from PBMC facilitating biological interpretation of CyTOF data.

**Fig 3.**
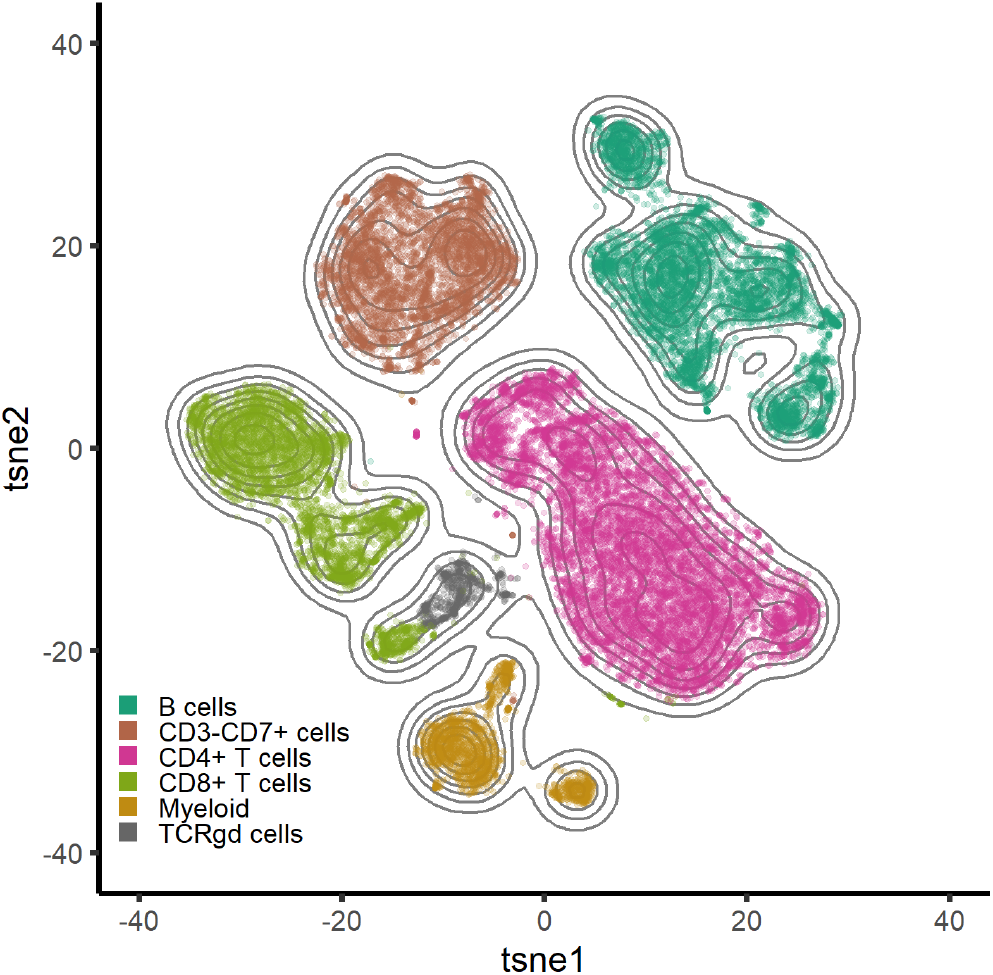
Recapitulation of cell types in the 12 PBMC samples using tSNE (n=30,000, perplexity=90, theta=0.4) colored by the combined cluster labels produced by FlowSOM.

### 3.3 Case study II: Cancer cell populations on HGSOC

To assess the performance of *Cyto* in enabling clinical interpretation we next performed an iterative analysis on a dataset of 15 clinical samples (Table S2) from HGSOC patients at diagnosis (primary), after neoadjuvant chemotherapy (interval) or at tumor progression. In this analysis *Cyto* also takes advantage of a detailed clinical metadata to assist variable association in the *results browser*.

Phenograph successfully detects main immune, stromal and tumoral cells (Figure 4A and Supplementary Figure S5). The immune compartment is the largest; we annotated the clusters to be CD8+ T-cells, CD8-CD3+ likely CD4+T-cells, and CD45+ T-cell marker negative likely Myeloid-lineage inflammatory cells. The stromal compartment is divided into CD90 positive and negative stromal cells, with the negative cells showing closer similarity to the tumor cells. The tumor compartment, identified as *Cluster-7* is characterized by high expression of EpCAM, MUC1, E-Cadherin and CA125, and low expression of pan-leucocyte marker CD45. Abundance difference (Figure 4B) show ascites samples (n=10) have more myeloid cells, and less tumor and stromal cells than solid tumor samples, while no apparent differences were observed on T-cell abundance. Interestingly, *Cluster-6* shows expression for stemness markers CD117 and CD44, the tumor markers CD125, HE4 and EpCAM, and is negative for the immune and stromal markers, presenting as a potential cancer stem cell population.

**Fig 4.**
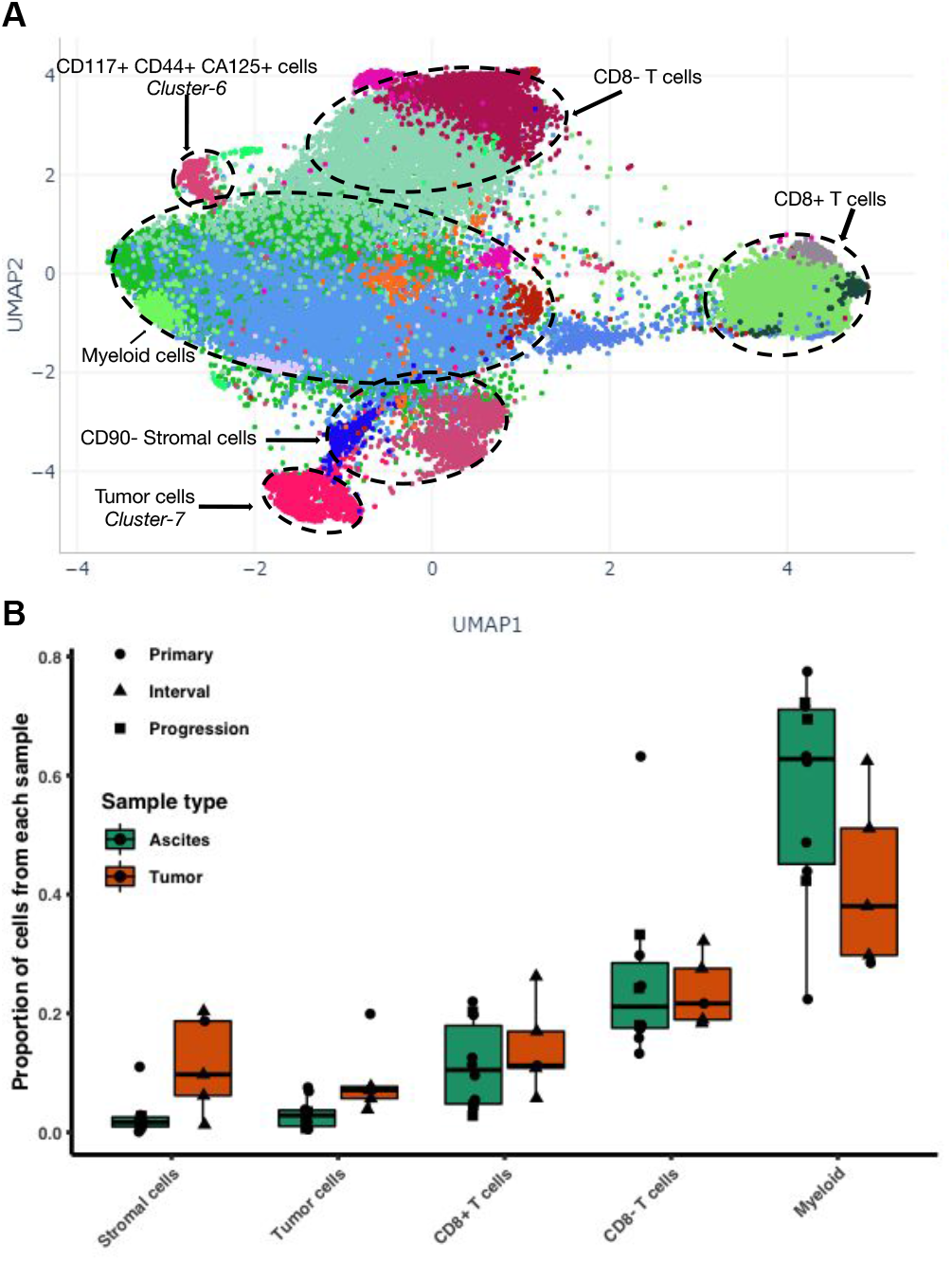
First iteration on high-grade serous ovarian cancer data. (A) Screenshot of all cells from 15 HGSOC samples from different therapy time-points and different tissue sites, Phenograph labels (colors) were computed with 300,000 cells randomly sampled and k=450. (B) Summary of proportions of cell types identified by Phenograph for each sample annotated with sample type and tissue type

A second iteration of Cyto analysis, in which we focused on the tumor cells (Figure 5 and Supplementary Figure S7), shows the integration of clinical annotations with a tumor subpopulation profiling analysis. The intermediate run that shows the detection of the tumor cells is shown in Figure S6. Minimum spanning tree (MST) representation of the detected clusters present distinct tumor population abundance in Primary, Interval, and Progression time of sampling. Furthermore, Cluster-6 on the MST shows higher Ki67 and more abundant in Primary and Interval samples. Cluster-2 shows highest E-Cadherin and is dominant in Interval samples, and progression samples have larger representation of Cluster-10, which are cells enriched for MUC1 and CD147, and are low on ERK1-2 signaling.

A Cyto visualization of Simpson’s diversity index highlights also that Progression samples have the lowest heterogeneity. Interestingly, collapsing the MST by time from sample to the next progression we see a clear enrichment of a stemness marker CD24 in samples with shorter time to progression.

**Fig 5.**
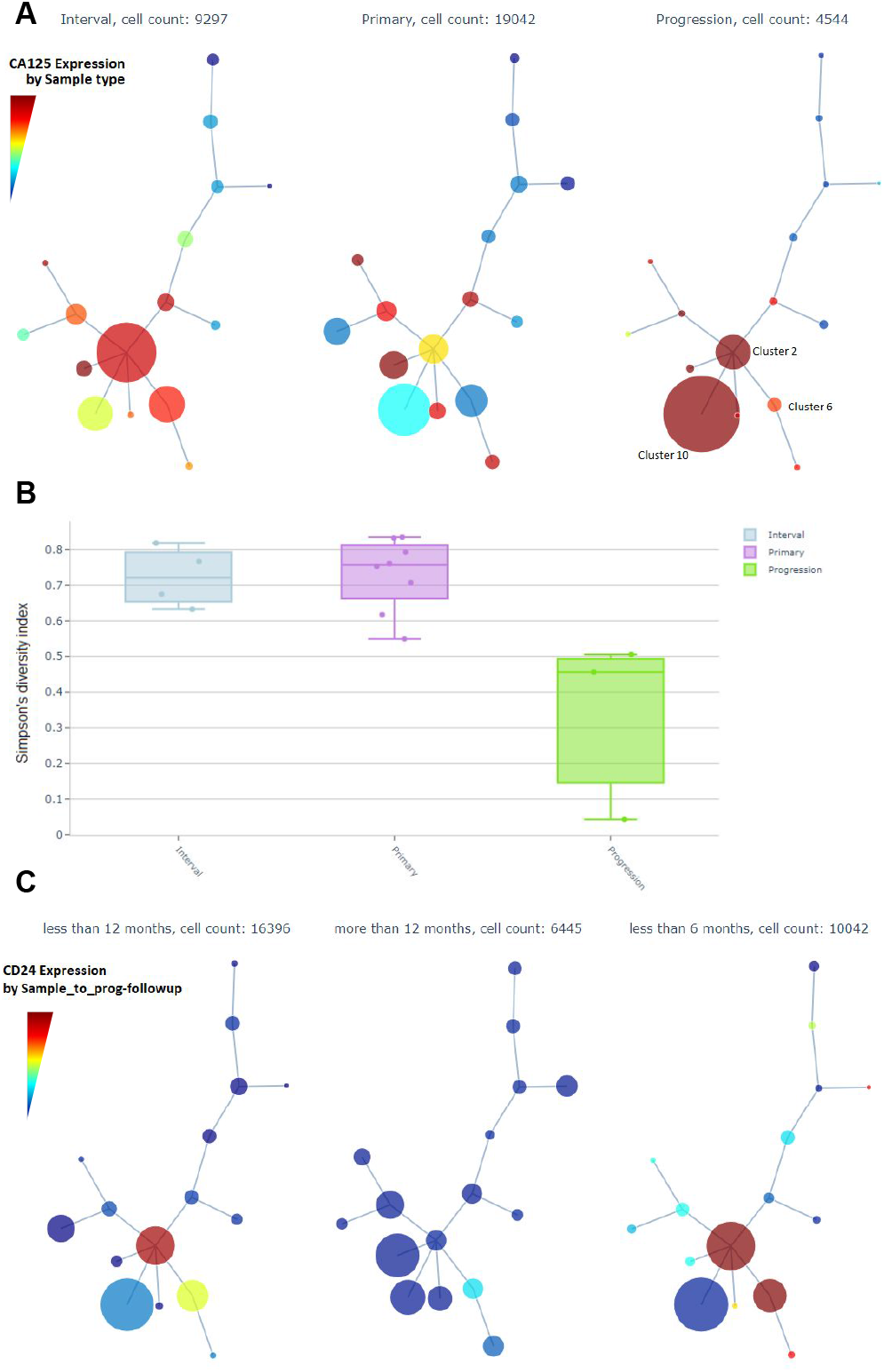
Screenshots of *Cyto* analysis of only tumor cell populations. (A) Minimum Spanning Trees (MST) by Sample time summarizes the expression of CA125(B) Simpson’s diversity index by Sample time. (C) CD24 expression across MST nodes grouped by time from sample to next progression.

## 4 Conclusion

Rapid advances in single-cell technologies produce larger and more complex data than ever before. Complex analyses increase the difficulty of reporting reproducible results, while accessibility to and usability of highly specialized tools drive the choice of algorithms in the analysis. A standard one-way analysis workflow is sufficient on low-dimensional data but a more exploratory research requires an iterative approach. We propose to level the usability of different tools and to ease reproducibility of analysis by integrating tools using a workflow paradigm design. First, by including popular cytometry methods as Anduril components available, less experienced bioinformaticians can easily build customized analysis workflows. Second, we present a generalized analysis pipeline that covers cytometry questions from detection of rare cells to differential abundance analysis, and from general sample profiling to deeper analysis of single cell populations. Third, by making this pipeline accessible as a Docker container with a user-friendly interface, non-bioinformaticians are able to perform complex single-cell analyses regardless of their experience level on software maintenance. Fourth, a side-effect of utilizing Docker for accessibilty includes the potential to run it remotely on a server.

To our knowledge *Cyto* is the first cytometry tool with a workflow paradigm design. Many R packages (Simpson, 2019; Spidlen. *et al.*, 2019; Finak, Greg *et al.*, 2014) have enabled compatibility with the popular flowCore package (Ellis *et al.*, 2019), and including them in our cytometry components allowes users to execute them as part of larger pipelines on computing clusters if necessary.

Additionally, this study demonstrates the key features of Cyto on a public, well-known dataset, as well as on a new independent cohort. Here we are able to identify and characterize cell population changes before and after chemotherapy, as well as at the time of progression. Ascites samples are valuable but underutilized due to the large number of non-tumor cells. Our analysis characterized the composition of there herein used ascites samples and the iterative analysis feature in Cyto enabled focusing on tumor cells without manual setting of thresholds for each sample. This allowed us to compare tumor cell phenotypes between clinical settings, suggesting that HGSOC tumors at relapse are characerized by higher heterogeneity and enriched stemness; an interesting hypothesis for further studies.

In summary, this work presents Cyto, which is an open-source, accessible and customizable cytometry analysis method that takes advantage of workflow engines and enables easy integration of existing tools. Cyto offers two levels for technical and non-technical users. Further, to our knowledge this study presents the first CyTOF experiments on comparison of chemotherapy naïve and heavily treated relapse samples from HGSOC.

## Acknowledgements

The authors thank all previous efforts in the cytometry field that enabled publicly available data and open access software, as well as Dr Daria Bulanova for helpful discussions. The authors also wish to thank sample donors for their invaluable contribution to cancer research.

for high-dimensional single-cell flow and mass cytometry data. Cytom. Part A, 89, 1084-1096.

## Supplementary Files

**Supplementary Material –** Additional methods and figures

**Supplementary File S1 –** Settings to reproduce figure 2

**Supplementary File S2 –** Settings to reproduce figure 3

**Supplementary File S3 –** Settings to reproduce figure 4A

**Supplementary File S4 –** Settings to reproduce figure 4B

## 1 Supplementary methods

### 1.1 Data acquisition of High-Grade Serous Ovarian Cancer

#### 1.1.1 Sample dissociation and preparation

Ovarian cancer primary cells were isolated from ascites and tumor tissues. Ascites was centrifuged at 3.0 G for 15 min, followed by gradient centrifugation with Histopaque-1077 to discard the contaminating blood cells from the sample. Tissues were cut in to approximately 1mm pieces and dissociated over night with 1:75 dilution of 10x Collagenase/hyaluronidase in warm DMEM-F12 media (Stem Cell technologies, Cambridge, UK). Cells were isolated by filtering the sample with 100μm and 70 μm meshes followed by Histopaque-1077 centrifugation to discard contaminating blood cells and cell debris.

#### 1.1.2 Antibody preparation for mass cytometry

The antibodies (Supplementary Table S2) were purchased already conjugated with metal isotopes from Fluidigm when available. Otherwise, purified carrier-free antibodies were purchased from other vendors (Biolegend, R&D System and BD Biosciences) and then conjugated with metal isotopes using the Maxpar antibody conjugation kit (Fluidigm) following the manufacturer’s instructions. In-house conjugated antibodies were quantified and diluted in PBS antibody stabilization solution (CANDOR Biosciences) to 0.1-0.4 mg/ml and stored at +4°C. CD166, CD133 and cleaved-PARP were purchased conjugated with fluorochromes and detected with anti-fluorochromes metal tagged antibodies (145Nd-PE, 176Yb-APC and 160Gd-FITC respectively) in a secondary staining step. Antibodies have been initially tested by flow cytometry followed by a titration at mass cytometer using cell lines to set the working concentration.

#### 1.1.3 Cell processing and antibody staining

The isolated cells were washed with 1x PBS, centrifuged, suspended in to warm DMEM-F12 medium and stained with 1 μM 103Rhodium-DNA Intercalator (Fluidigm). Unstained cells were acquired as control sample to detect the background signals. After 15 min incubation at 37°C, cells were washed with Cell Staining Medium (CSM) [PBS, 0.5% BSA (Sigma Aldrich), 0.02 % NaN3 (Sigma Aldrich)] and fixed with 1.6% paraformaldehyde (Electron Microscopy Sciences) for 10 min at room temperature. Samples were washed twice with CSM and shipped at +4°C to the Istituto Superiore di Sanità, Roma, Italy. Upon arrival the cells were counted and around 2-3 million were pelleted, washed with CSM and incubated with Fc-blocker (Biolegend) for 10 minutes at RT to counteract the antibody binding to FC-receptors. Cellular staining with antibodies was performed according to the manufacturer’s protocol (Fluidigm) consisting of several staining steps interspersed with permeabilization treatments moving from gentlest to strongest conditions. In the first step, cells were resuspended in 100 μl of a mix of CSM and metal-conjugated antibodies specific for surface antigens, incubated for 30 minutes at RT and washed twice in CSM. In a second step the cells were permeabilized with 1ml of CSM supplemented with 0.3% Saponin (Sigma Aldrich) (CSM-S) for 30 minutes at +4°C and then stained with an antibody cocktail specific for intracellular antigens, for 45 minutes at RT and washed twice with CSM-S. The third step consisted in a stronger permeabilization of the cell pellet with 1 ml of ice cold methanol (Sigma Aldrich) per 0.5×10^6^ cells for 10 minutes at 4°C. Cells were then washed twice in CSM and stained with 100 μl of a further antibody mix toward phosphoproteins and transcription factors, for 60 minutes at RT in CSM. After washing with CSM, the cells were stained with 125 nM 191/193Iridium-DNA Intercalator (Fluidigm), in PBS/PFA 1.6% for 20 minutes at RT (or overnight at 4), for cell events recognition during data acquisition, and then washed twice with CSM and once with MilliQ water.

#### 1.1.4 CyTOF assay and data preprocessing

Before acquisition, cells were counted and diluted at 2×10^5^ cells/ml in MilliQ water with 1/10 of volume of EQ^TM^ Four Element Calibration Beads (Fluidigm) and filtered through a 35μm nylon mesh before acquisition. Data from each sample were pre-processed with CyTOF software version 6.7.1014 to normalize signals and minimize instrument performance variation during acquisition (lower convolution threshold of 200, event length between 10 and 75 and with a rate of 500 cells/sec). FCS files were processed with FlowJo software (FlowJo LLC) to export bead-normalized single-viable cells based on gating performed on cell length and DNA intercalators signals (191/193Iridium and 103Rhodium). Because each sample was processed at the time of acquisition to conserve signal quality, the header of the raw FCS files were matched in R.

## 2 Supplementary tables

**Table S1.** Methods and tools available within the Anduril pipeline integrated in *Cyto.*

| *Category* | *Method/Tool* |
| --- | --- |
| *Preprocessing* | Density-biased sampling |
|  | Random down sampling |
|  | Data tranformation |
|  | Sample-wise normalization |
| *Clustering* | Phenograph |
|  | FlowSOM |
|  | FlowMeans |
|  | XShift |
|  | K-means |
| *2D Embedding* | tSNE |
|  | UMAP |
| *Build Data Dashboard* | MDS |
|  | NRS |
|  | Minimum Spanning trees |

**Table S3.**
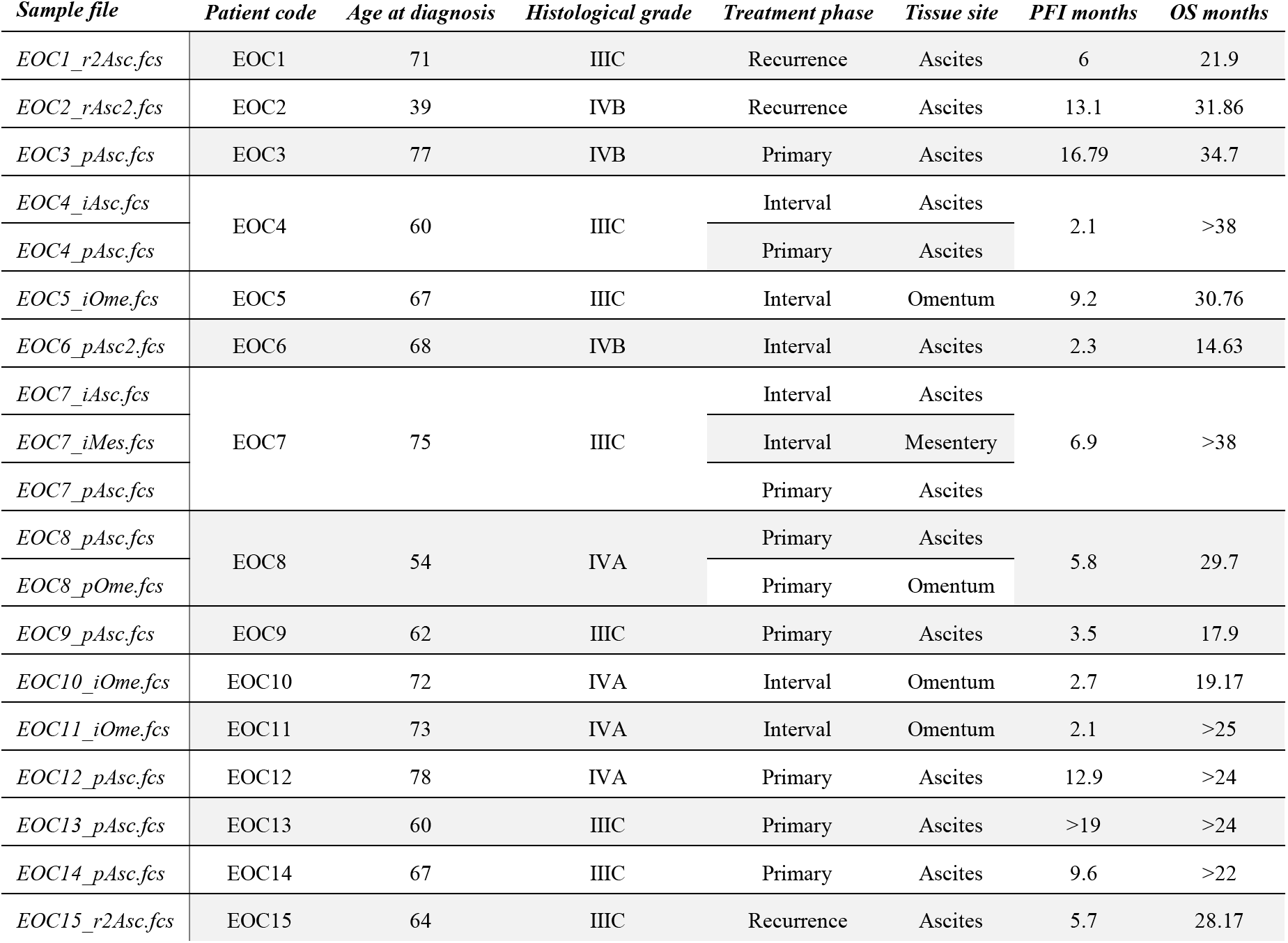
Sample cohort included in this study. Age, stage, tissue, number of patients, number of total cells acquired from each, survival, treatment.

| *Sample file* | *Patient code* | *Age at diagnosis* | *Histological grade* | *Treatment phase* | *Tissue site* | *PFI months* | *OS months* |
| --- | --- | --- | --- | --- | --- | --- | --- |
| *EOC1_r2Asc.fcs* | EOC1 | 71 | IIIC | Recurrence | Ascites | 6 | 21.9 |
| *EOC2_rAsc2.fcs* | EOC2 | 39 | IVB | Recurrence | Ascites | 13.1 | 31.86 |
| *EOC3_pAsc.fcs* | EOC3 | 77 | IVB | Primary | Ascites | 16.79 | 34.7 |
| *EOC4_iAsc.fcs* | EOC4 | 60 | IIIC | Interval | Ascites | 2.1 | >38 |
| *EOC4_pAsc.fcs* |  |  |  | Primary | Ascites |  |  |
| *EOC5_iOme.fcs* | EOC5 | 67 | IIIC | Interval | Omentum | 9.2 | 30.76 |
| *EOC6_pAsc2.fcs* | EOC6 | 68 | IVB | Interval | Ascites | 2.3 | 14.63 |
| *EOC7_iAsc.fcs* | EOC7 | 75 | IIIC | Interval | Ascites | 6.9 | >38 |
| *EOC7_iMes.fcs* |  |  |  | Interval | Mesentery |  |  |
| *EOC7_pAsc.fcs* |  |  |  | Primary | Ascites |  |  |
| *EOC8_pAsc.fcs* | EOC8 | 54 | IVA | Primary | Ascites | 5.8 | 29.7 |
| *EOC8_pOme.fcs* |  |  |  | Primary | Omentum |  |  |
| *EOC9_pAsc.fcs* | EOC9 | 62 | IIIC | Primary | Ascites | 3.5 | 17.9 |
| *EOC10_iOme.fcs* | EOC10 | 72 | IVA | Interval | Omentum | 2.7 | 19.17 |
| *EOC11_iOme.fcs* | EOC11 | 73 | IVA | Interval | Omentum | 2.1 | >25 |
| *EOC12_pAsc.fcs* | EOC12 | 78 | IVA | Primary | Ascites | 12.9 | >24 |
| *EOC13_pAsc.fcs* | EOC13 | 60 | IIIC | Primary | Ascites | >19 | >24 |
| *EOC14_pAsc.fcs* | EOC14 | 67 | IIIC | Primary | Ascites | 9.6 | >22 |
| *EOC15_r2Asc.fcs* | EOC15 | 64 | IIIC | Recurrence | Ascites | 5.7 | 28.17 |

**Table S2.** Antibodies used for in-house CyTOF data.

| *Cat No* | *Metal Tag* | *Target* | *Clone* | *Vendor* | *Final Concentration (µg/100 µl)* | *Dilution* |
| --- | --- | --- | --- | --- | --- | --- |
| 3141006B | 141Pr | EpCAM | 9C4 | Fluidigm | N.A | 1:50 |
| 3143001B | 143Nd | CD117 | 104D2 | Fluidigm | N.A | 1:50 |
| MAB56091 | 144Nd | CA125 | 986811 | R&D | 1.2 | - |
| 559263/3145006B | 145Nd | CD166-PE | 3A6/PE001 | BD/Fluidigm | N.A | 1:30/1:50 |
| 3147015B | 147Sm | ALDH | 44/ALDH | Fluidigm | N.A | 1:50 |
| 304045 | 149Sm | CD45 | HI30 | Biolegend | 0,6 | - |
| 561469 | 151Eu | Sox2 | O30-678 | BD | 1.2 | - |
| 3152005A | 152Sm | pAkt [S473] | D9E | Fluidigm | N.A | 1:50 |
| 3153021B | 153Eu | CD44s Pan Specific | 691534 | Fluidigm | N.A | 1:75 |
| 3154003B | 154Sm | CD3 | UCHT1 | Fluidigm | N.A | 1:75 |
| 355602 | 155Gd | MUC1 | 16A | Biolegend | 0.7 | - |
| 3156022B | 156Gd | CD147 | HIM6 | Fluidigm | N.A | 1:75 |
| 3158021A | 158Gd | E-Cadherin | 24E10 | Fluidigm | N.A | 1:50 |
| 3159029B | 159Tb | PD-L1 | 29E.2A3 | Fluidigm | N.A | 1:50 |
| 558576/3160011B | 160Gd | Cleaved PARP-FITC | F21-852/FIT22 | BD/Fluidigm | N.A | 1:30-1:50 |
| 3161009B | 161Dy | CD90 | 5E10 | Fluidigm | N.A | 1:75 |
| 3162015B | 162Dy | CD8a | RPA-T8 | Fluidigm | N.A | 1:75 |
| MAB6274 | 164Dy | HE4 | 676013 | R&D | 0.9 | - |
| 350802 | 165Ho | N-Cadherin | 8C11 | Biolegend | 0.9 | - |
| 550314 | 166Er | CD146 | P1H12 | BD | 1.2 | - |
| 3171010A | 167Er | pERK 1/2 [T202/Y204] | D13.14.4E | Fluidigm | N.A | 1:50 |
| 3168007B | 168Er | Ki-67 | B56 | Fluidigm | N.A | 1:50 |
| 3169004B | 169Tm | CD24 | ML5 | Fluidigm | N.A | 1:50 |
| 3172014B | 172Yb | PD-L2 | 24F.10C12 | Fluidigm | N.A | 1:50 |
| 3174020B | 174Yb | PD-1 | EH12.2H7 | Fluidigm | N.A | 1:50 |
| 3175009A | 175Lu | pS6 [S235/S236] | N7-548 | Fluidigm | N.A | 1:50 |
| 130-090-826/3176007B | 176Yb | CD133-APC | AC133/APC003 | Miltenyi/Fluidigm | N.A | 1:30-1:50 |

## 3 Supplementary figures

**Figure S1.**
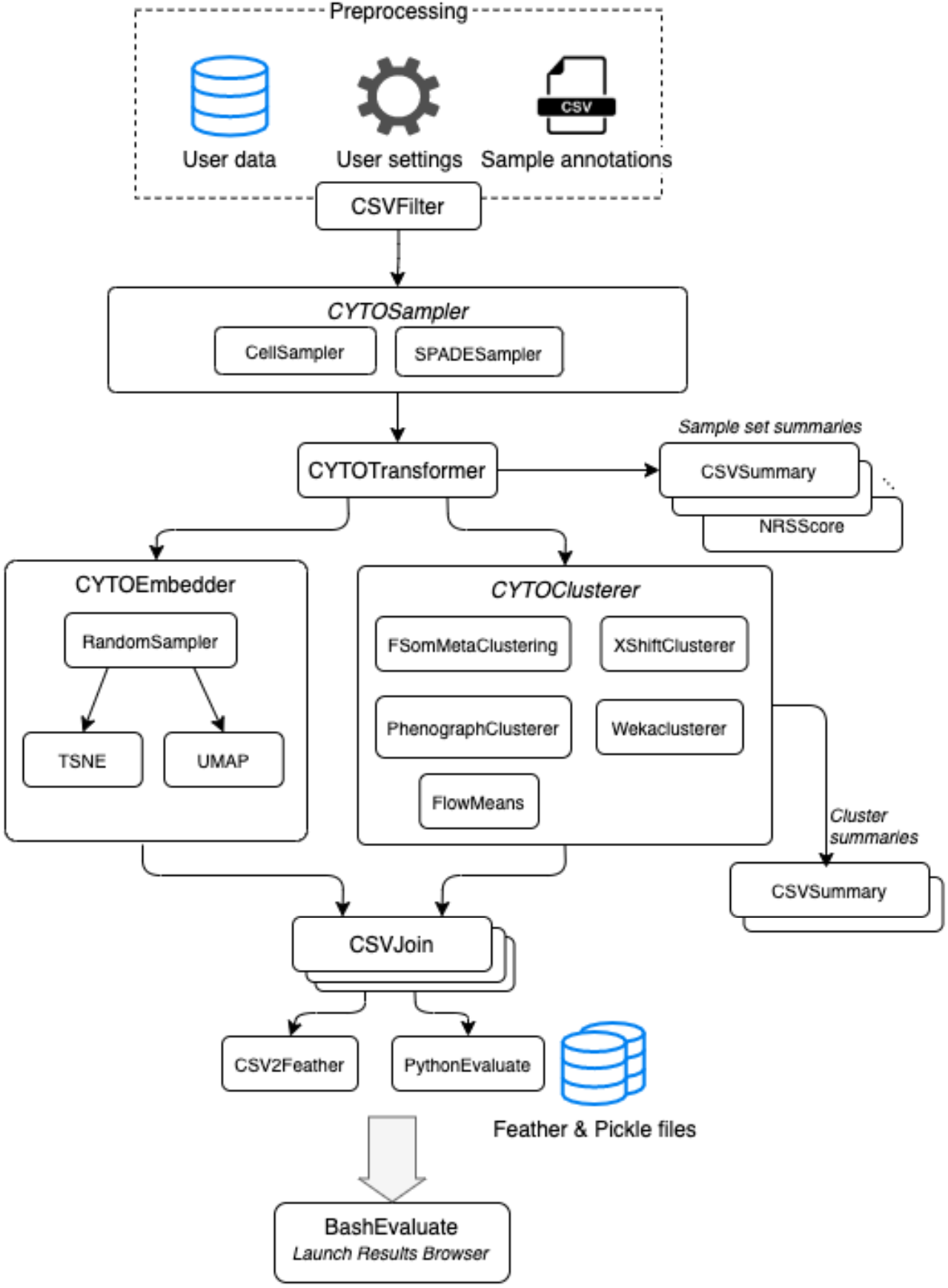
Detailed workflow of the Anduril workflow used in *Cyto.*

**Figure S2.**
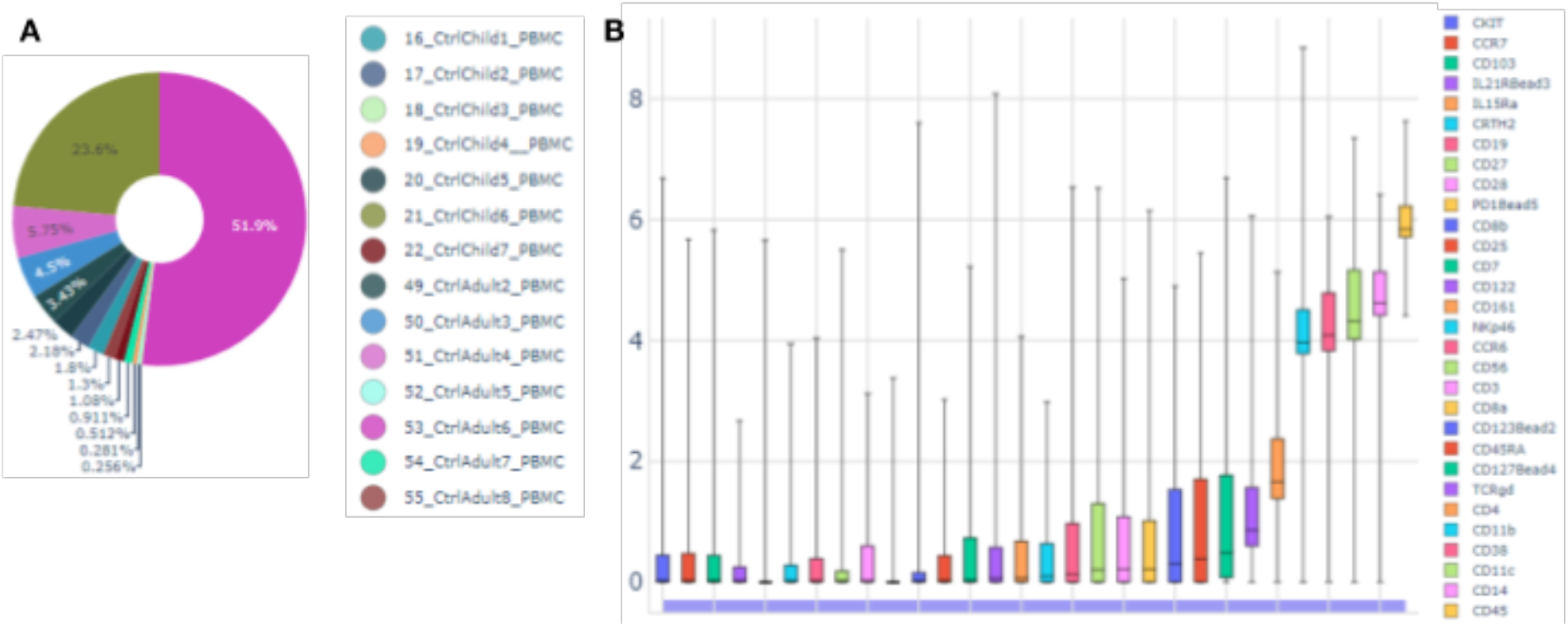
(A) Sample representation within the dominant cluster from the outlier sample 53_CtrlAdul6_PBMC. (B) Marker signal distribution within this cluster. The highest expressing markers are CD14, CD11c, CD38, and CD11b.

**Figure S3.**
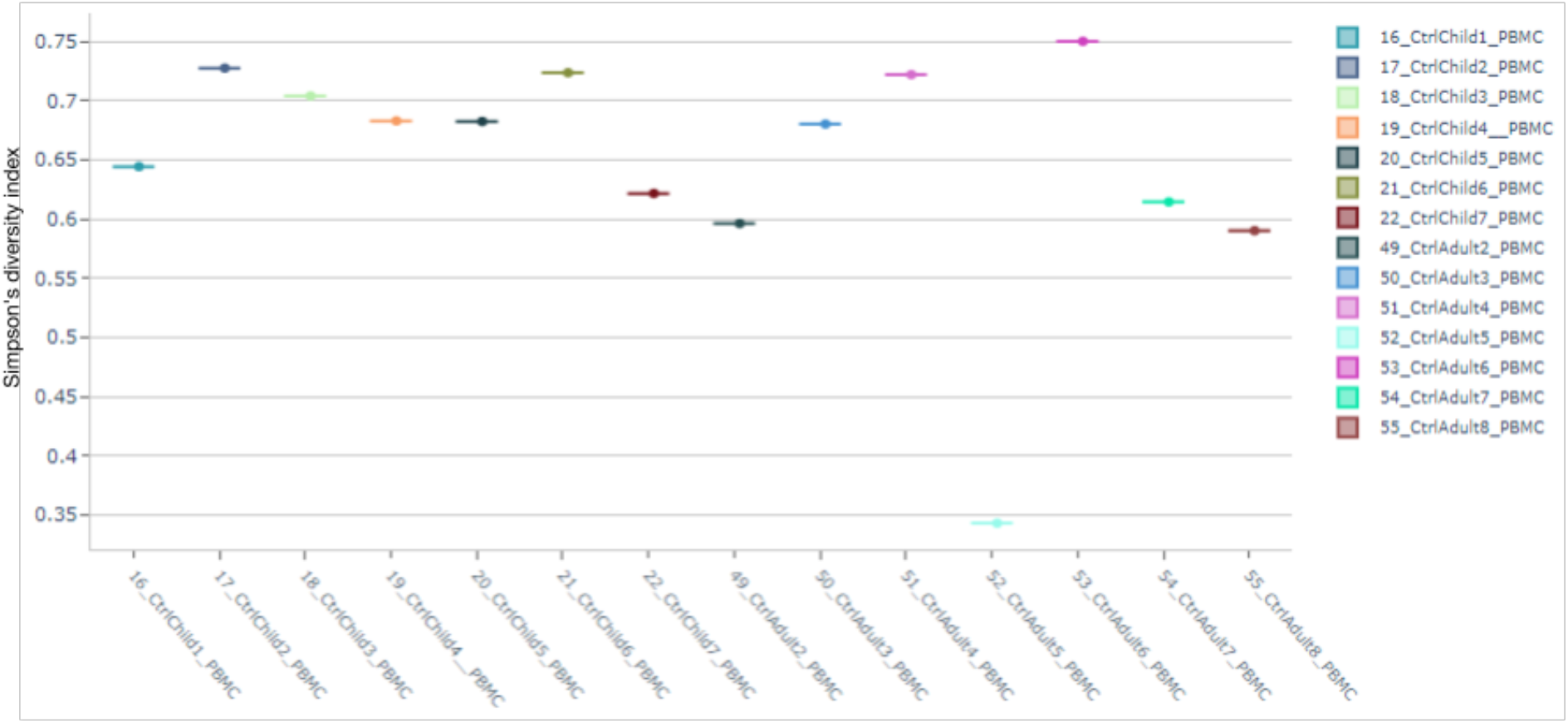
Simpson’s diversity index identifies sample 52_CtrlAdult5_PBMC (in Cyan) as an outlier based on the number of clusters represented within the sample and the relative abundance of each of the clusters.

**Figure S4.**
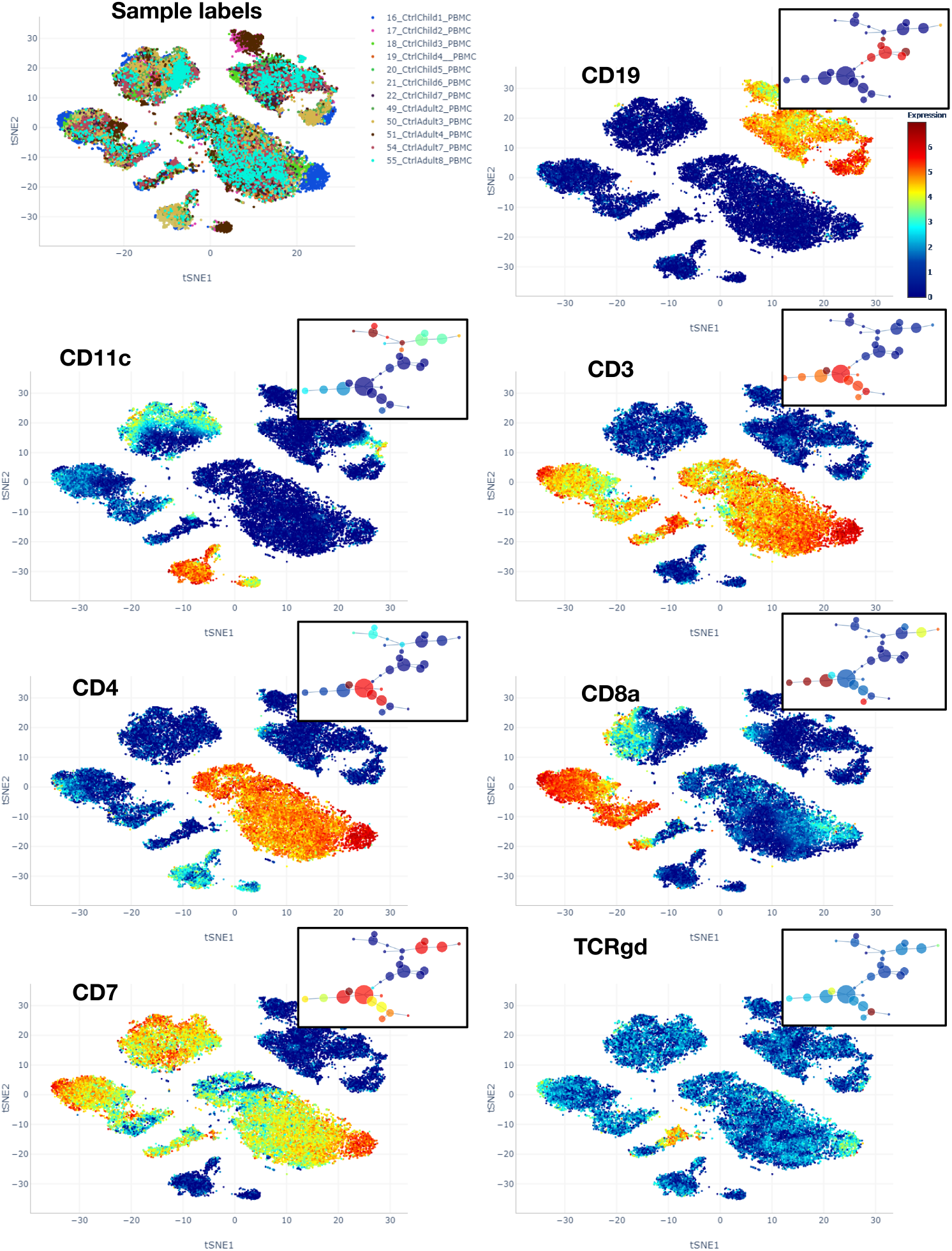
Expression profiles of the PBMC cell types without the two outlier samples. The sample labels indicate that a batch effect is not dominant. Visualization of the dataset as tSNE or as MST shows the expression and relationships of the populations shown in Figure 3.

**Figure S5.**
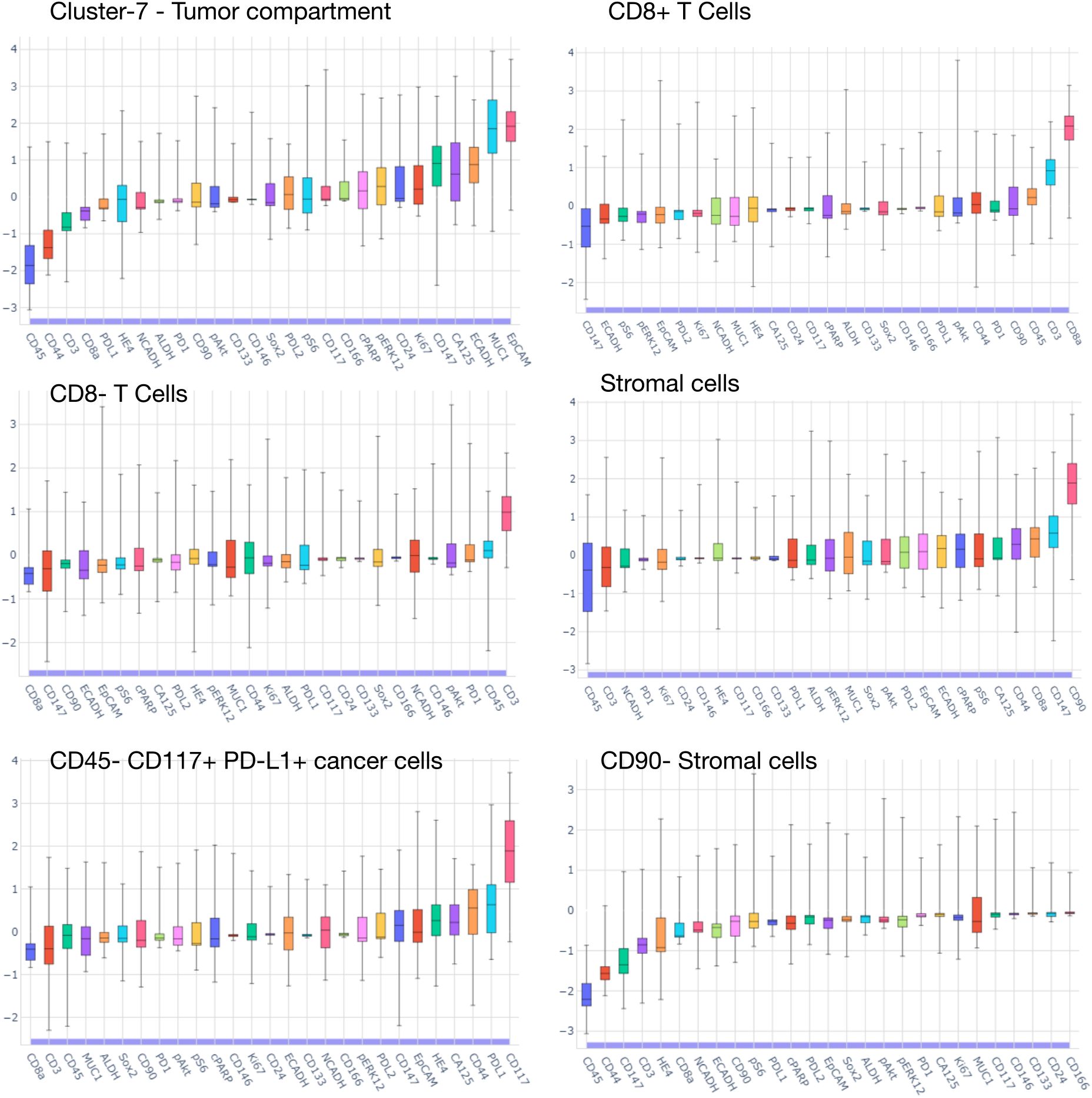
Expression distribution of each marker within the cell clusters selected in Figure 4. The lasso selection tool in the results browser allows to explore the expression profiles interactively.

**Figure S6.**
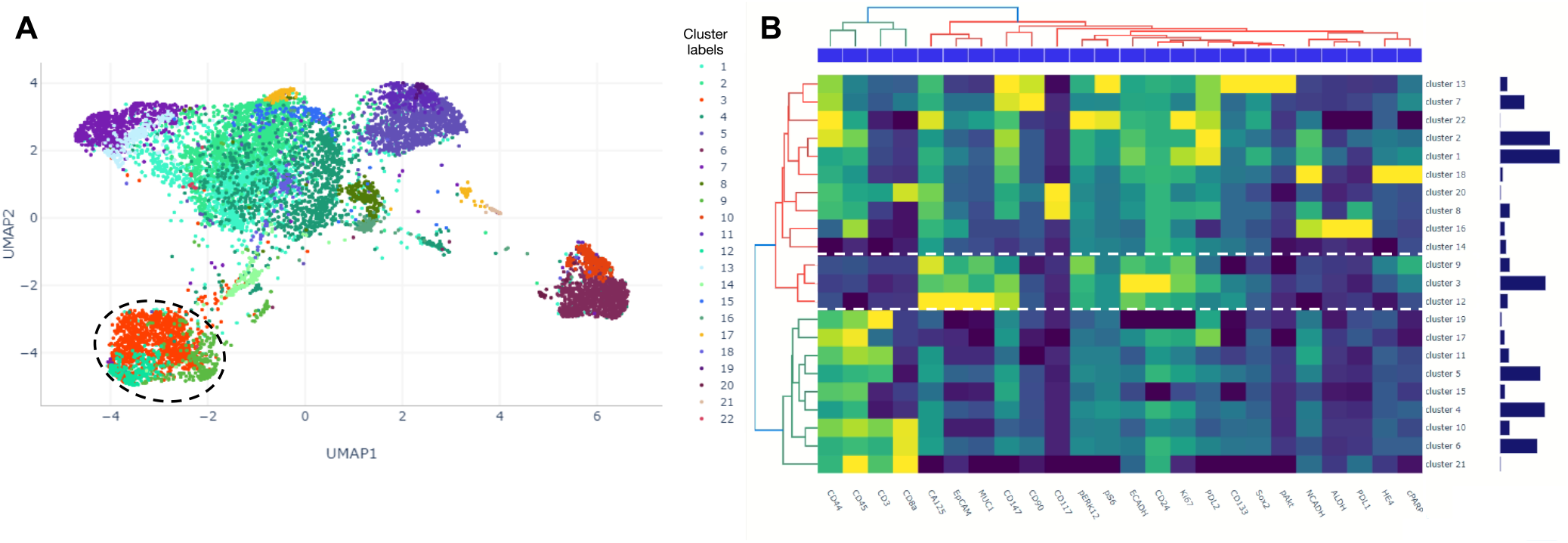
Identification of the tumor compartment using density-based sampling option. (A) UMAP of the density-based sampled dataset with the tumor cells highlighted with a dashed line. (B) Expression profile of each cluster. Tumor cells are highlighted with a dashed line and further filtered for the tumor cell analysis.

**Figure S7.**
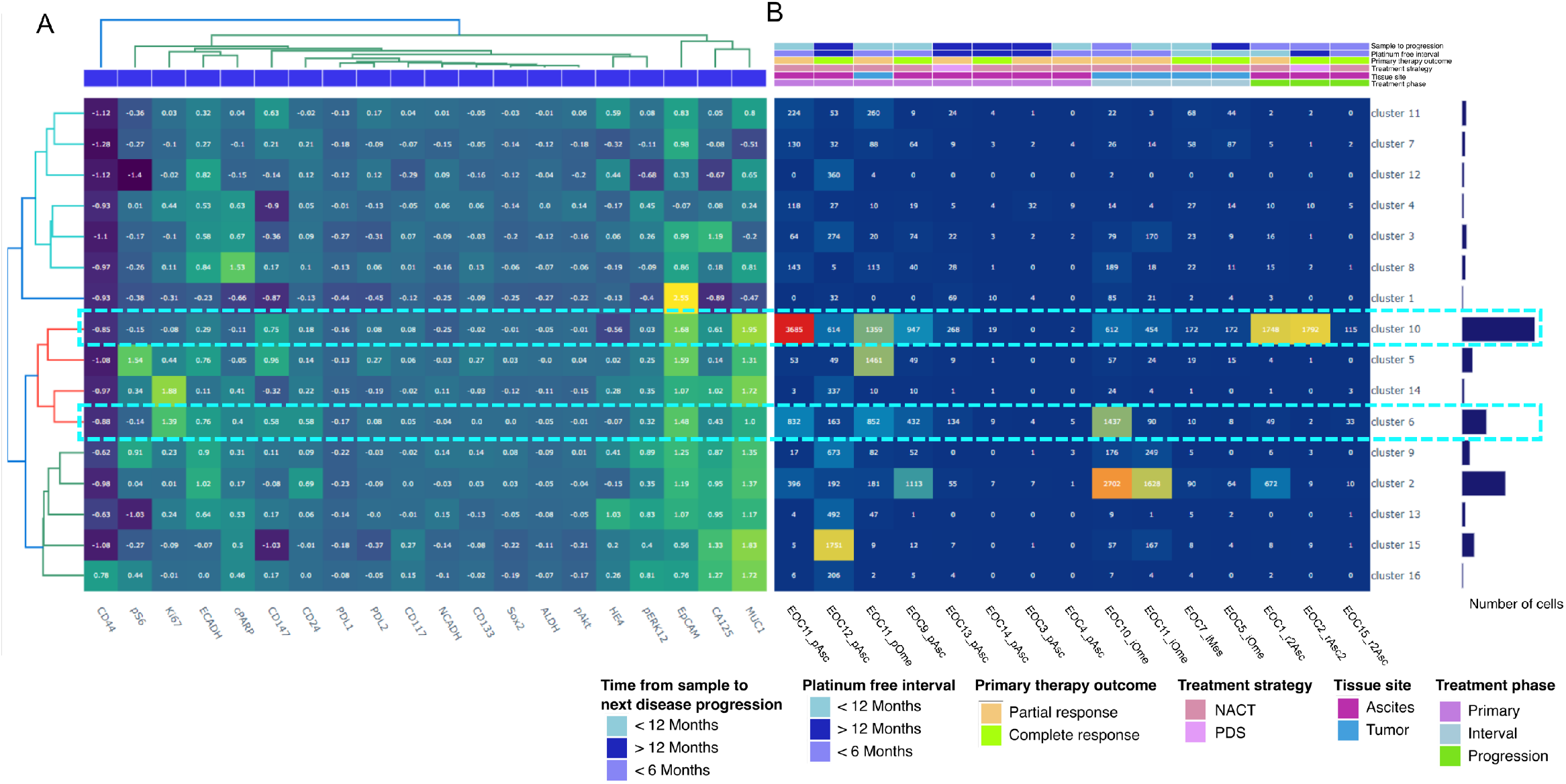
Tumor cell clustering with focus on Cluster-10 and Cluster-6 (highlighted in cyan) (A) Marker expression and hierarchical clustering of the subpopulations. (B) Population abundance across the samples.

## References

Almeida, J.S. (2010) Computational ecosystems for data-driven medical genomics. Genome Med., 2, 67.

Amir, E.A.D. et al. (2013) ViSNE enables visualization of high dimensional single-cell data and reveals phenotypic heterogeneity of leukemia. Nat. Biotechnol., 31, 545–552.

Angerer, P. et al. (2016) *destiny:* diffusion maps for large-scale single-cell data in R. Bioinformatics, 32, 1241–1243.

Brodin, P. (2018) The biology of the cell - insights from mass cytometry. FEBS J.

Cervera, A. et al. (2019) Anduril 2: upgraded large-scale data integration framework. Bioinformatics.

Chen, H. et al. (2016) Cytofkit: A Bioconductor Package for an Integrated Mass Cytometry Data Analysis Pipeline. PLOS Comput. Biol., 12, e1005112.

Dix, A. and Ellis, G. (1998) Starting Simple: Adding Value to Static Visualisation Through Simple interaction. In, Proceedings of the working conference on Advanced visual interfaces.

Ellis, B. et al. (2019) flowCore: flowCore: Basic structures for flow cytometry data.

Finak, Greg et al. (2014) OpenCyto: An Open Source Infrastructure for Scalable, Robust, Reproducible, and Automated, End-to-End Flow Cytometry Data Analysis. PLoS Comput. Biol., 10, e1003806.

Galli, E. et al. (2019) The end of omics? High dimensional single cell analysis in precision medicine. Eur. J. Immunol.

Höllt, T. et al. (2016) Cytosplore: Interactive Immune Cell Phenotyping for Large Single-Cell Datasets. Comput. Graph. Forum, 35, 171–180.

Kotecha, N. et al. (2010) Web-Based Analysis and Publication of Flow Cytometry Experiments. Curr. Protoc. Cytom., 53, 10.17.1-10.17.24.

Van Der Maaten, L. and Hinton, G. (2008) Visualizing Data using t-SNE.

Nowicka, M. et al. (2017) CyTOF workflow: differential discovery in high-throughput high-dimensional cytometry datasets. F1000Research, 6, 748.

Qiu, P. et al. (2011) Extracting a cellular hierarchy from high-dimensional cytometry data with SPADE. Nat. Biotechnol., 29, 886–893.

Qiu, P. (2017) Toward deterministic and semiautomated SPADE analysis. Cytom. Part A, 91, 281–289.

Simpson, S. (2019) flowAssist: User friendly manipulation and analysis of flow cytometry data.

Spidlen., J. et al. (2019) flowUtils: Utilities for flow cytometry.

van Unen, V. et al. (2017) Visual analysis of mass cytometry data by hierarchical stochastic neighbour embedding reveals rare cell types. Nat. Commun., 8, 1740.

Van Unen, V. et al. (2016) Mass Cytometry of the Human Mucosal Immune System Identifies Tissue-and Disease-Associated Immune Subsets.

Weber, L.M. and Robinson, M.D. (2016) Comparison of clustering methods for high-dimensional single-cell flow and mass cytometry data. Cytom. Part A, 89, 1084–1096.

